# Age dependence of modern clinical risk groups for localized prostate cancer – a population-based study

**DOI:** 10.1101/648519

**Authors:** Minh-Phuong Huynh-Le, Tor Åge Myklebust, Christine H. Feng, Roshan Karunamuni, Tom Børge Johannesen, Anders M. Dale, Ole A. Andreassen, Tyler M. Seibert

## Abstract

**Background:** Optimal prostate cancer (PCa) screening strategies will focus on men most likely to have potentially-lethal, localized disease. Age-specific incidence rates (ASIRs) for clinical risk groups could guide risk-stratified screening.

**Objective:** Determine ASIRs and proportions of PCa diagnoses in Norway for modern risk-group and Gleason score categories.

**Design, Setting, and Participants:** All men diagnosed with PCa in Norway in 2014-2017 (n=20,356).

**Outcome Measurements and Statistical Analysis:** Patients were assigned to clinical risk groups: low, favorable-intermediate, unfavorable-intermediate, high, regional, and metastatic, using Gleason score and clinical stage. Associations were assessed between age and (1) Gleason score (including Gleason 3+4 and 4+3) and (2) PCa risk group. Risk-group ASIRs were calculated by multiplying the overall Norwegian ASIR by the proportions observed for each category.

**Results:** Older age was significantly associated with higher Gleason score and more advanced disease. For example, among men aged 55-59, 65-69, 75-79, and 85-89 years, the percentage with Gleason 8-10 disease was 16.5%, 23.4%, 37.2%, and 59.9%, respectively (p<0.001); the percentage with at least high-risk disease was 29.3%, 39.1%, 60.4%, and 90.6%, respectively. Corresponding percentages for low-risk PCa were 24.0%, 17.9%, 10.2%, and 4.1% (p<0.001). The respective maximum ASIRs (per 100,000 men) for low-risk, favorable-intermediate-risk, unfavorable-intermediate-risk, high-risk, regional, and metastatic disease were: 157.1, 183.8, 194.8, 408.3, 172.3, and 330.0; incidence for low-risk and favorable-intermediate-risk PCa peaked before age 70, while more advanced categories peaked after 70. At age 75-79 years, the ASIR of high-risk disease was approximately 6 times greater than at 55-59 years.

**Conclusions:** Risk of clinically-significant, localized PCa increases with age. Healthy older men may be among those most likely to benefit from PCa screening.

## Introduction

Most efforts regarding prostate cancer (PCa) screening now seek to identify clinically-significant and potentially-lethal cases requiring treatment, while avoiding overdiagnosis of more indolent, lower-risk cases eligible for active surveillance^1,2^. Disease aggressiveness is accounted for in clinical decision-making with widely-used stratification schemes for risk of metastasis or death^3,4^. Men diagnosed with intact PCa may be followed with active surveillance^5^ or treated with surgery, radiotherapy, androgen deprivation therapy, or a combination thereof^3^. Understanding the age-specific incidence of modern clinical risk groups (defined below) could greatly inform effective screening strategies aimed to detect potentially-lethal, localized, PCa.

Prior studies have suggested associations between older age and both higher Gleason score^6,7^ and higher-risk disease^8^. These associations have important implications for pre-test probability for screening and diagnostic tests for clinically-significant PCa. Modern clinical risk groups (hereafter referred to as “risk groups”) include a distinction between unfavorable- and favorable-intermediate-risk cancer^9^. Unfavorable-intermediate-risk and high-risk disease have similar rates of developing distant metastases and of prostate-cancer-specific mortality^10^. This differentiation has been incorporated into clinical management^3^ and staging guidelines^11^. The difference between Gleason 4+3 and 3+4 is also included in modern guidelines^3,9–11^, but age associations have not been reported for this distinction. Additionally, age-specific incidence rates (ASIRs) for PCa are unknown for risk groups in current use.

Here, we used Norwegian population data to assess associations between age and PCa risk groups. We also estimated ASIRs in Norway for each risk group. Using modern strata for Gleason score and clinical risk group, we hypothesized that older men are more likely to have more advanced PCa.

## Methods

### Patient Population

We identified all men with PCa diagnosed from 2014-2017 from the Cancer Registry of Norway. Overall, the registry has been reported to have over 99% validity and completeness for PCa reporting^12,13^. Information potentially available included: age at diagnosis, clinical TNM stage, and Gleason score (including primary and secondary Gleason score, allowing distinction of Gleason 3+4 from 4+3, and favorable-from unfavorable-intermediate-risk^14^). These data are reported directly from clinicians/hospitals on digital reporting forms. Patients were placed into five-year age groups for analysis.

### Risk Stratification

PCa risk stratification was based on National Comprehensive Cancer Network (NCCN) guidelines^3^. Very-low-risk and low-risk case are managed similarly, so these were combined into low-risk PCa here (Gleason score ≤6, clinical stage T1-T2, prostate-specific antigen [PSA]<10 ng/mL). Clinical stage T2 was not specified in the registry as T2a, T2b, or T2c; we conservatively assumed that all T2 cases were T2a. High-risk and very-high-risk cases are managed similarly; these were combined into high-risk PCa (clinical stage ≥T3a or Gleason score ≥8 or PSA>20 ng/mL). Regional and metastatic cases had diagnosis of regional lymph node or distant metastatic disease, respectively. Intermediate-risk cases were all that did not fit the above definitions. Favorable-intermediate-risk cases had <50% positive biopsy cores and only one of the following: (1) Gleason score 3+4 with PSA<10 ng/mL or (2) Gleason score 3+3 with PSA 10-20 ng/mL. Unfavorable-intermediate-risk cases had any of the following: (1) Gleason score 4+3, (2) Gleason score 3+4 with PSA 10-20 ng/mL, (3) Gleason score 3+3 with PSA 10-20 ng/mL, or (4) ≥50% positive biopsy cores.

### Statistical Analyses

We used *R* for all analyses and chi-squared tests (two-tailed alpha=0.05) for all associations^15^. We tested for association between age and Gleason score at diagnosis, with separation of Gleason 7 disease into Gleason 3+4 and Gleason 4+3. Among those with low-grade (e.g., Gleason 6) disease, we tested for association between age and higher risk group due to PSA≥10 ng/mL and/or high clinical stage (T3-T4 or nodal or distant metastasis), as these are often ineligible for active surveillance, despite the low Gleason score^3^.

An association between age and PCa risk groups was also tested; this included separation of the intermediate-risk group category into favorable and unfavorable disease. We further tested for association between age and clinically-significant PCa (i.e., less appropriate for active surveillance^3^), which was defined two ways: first, all cases of at least intermediate-risk, and second, all cases of at least unfavorable-intermediate-risk. The above Chi-squared tests were repeated, while limiting analyses to only cases diagnosed between ages 50-74^3,16^.

Finally, we calculated PCa ASIRs for clinical risk groups. ASIRs for any PCa in Norway in 2015 (per 100,000 males) were obtained from NORDCAN, a comprehensive database of cancer statistics for Nordic countries^17,18^. This overall ASIR was multiplied by the proportion of cases meeting criteria for high-risk PCa to estimate the ASIR of high-risk PCa, specifically. This was repeated to obtain ASIRs for each risk group.

## Results

Of the 20,356 men diagnosed with PCa in Norway, 18,665 (91.7%) had Gleason score available. 17,920 patients (88.0%) could be categorized as low risk or not (i.e., intermediate risk or higher); 14,303 (70.3%) had sufficient available clinical data to assign a modern clinical risk group (including favorable/unfavorable-intermediate-risk). Lack of PSA and/or T stage data precluded assignment of a risk group for the remaining cases.

Older age at diagnosis was significantly associated with a higher Gleason score. **Figure 1** shows all age-stratified proportions of cases by Gleason score (proportions are listed in **eTable 1)**. The percentage of men with Gleason 8-10 disease among men aged 55-59, 65-69, 75-79, and 85-89 years was 16.5%, 23.4%, 37.2%, and 59.9%, respectively (p<0.001). Of those with Gleason 7 disease, older patients were more likely to be diagnosed with Gleason 4+3 compared to Gleason 3+4 disease. The percentage of Gleason 3+4 disease among men aged 55-59, 65-69, 75-79, and 85-89 years was 35.7%, 32.5%, 24.0%, and 12.4%, respectively, while the percentage of Gleason 4+3 disease across the same age groups was 14.0%, 17.3%, 19.3%, and 14.3%, respectively (p<0.001). When evaluating only those with Gleason 6, older men were more likely to be diagnosed with disease ineligible for active surveillance (**Figure 2**). The percentage of men aged 55-59, 65-69, 75-79, and 85-89 years with Gleason 6 disease and one or more of these risk factors (PSA ≥10 ng/mL, clinical stage ≥T3a, or nodal/metastatic disease) was: 19.5%, 28.7%, 42.9%, and 58.0%, respectively (p<0.001). Similar trends were seen for those men with Gleason 3+4 disease, and when men with Gleason 6 and 3+4 disease were combined (**eFigure 1**).

**Table 1.** Prostate cancer age-specific incidence rates (ASIRs) in Norway per 100,000 men by clinical risk group stratification. Intermediate-risk prostate cancer is subdivided into favorable and unfavorable risk.

| **Age**<br>**(years)** | **ASIRs (per 100,000 men) by prostate cancer risk group** |  |  |  |  |  |
| --- | --- | --- | --- | --- | --- | --- |
|  | **Low-risk** | **Favorable**<br>**intermediate-**<br>**risk** | **Unfavorable**<br>**Intermediate-**<br>**risk** | **High-risk** | **Regional** | **Metastatic** |
| <50 | 10.4 | 9.3 | 10.0 | 4.2 | 1.3 | 1.7 |
| 50-54 | 24.1 | 20.5 | 24.8 | 15.0 | 2.7 | 3.8 |
| 55-59 | 72.2 | 68.3 | 72.2 | 61.6 | 13.2 | 13.6 |
| 60-64 | 115.1 | 126.2 | 139.5 | 145.2 | 29.5 | 28.3 |
| 65-69 | 157.1 | 183.8 | 194.6 | 245.6 | 50.2 | 47.0 |
| 70-74 | 127.0 | 173 | 194.8 | 328.7 | 61.4 | 72.5 |
| 75-79 | 106.0 | 144.6 | 159.9 | 408.3 | 85.1 | 132.7 |
| 80-84 | 51.5 | 63.3 | 80.2 | 346.9 | 124.9 | 166.3 |
| 85+ | 24.0 | 8.8 | 14.2 | 222.3 | 172.3 | 330.0 |

**Figure 1.**
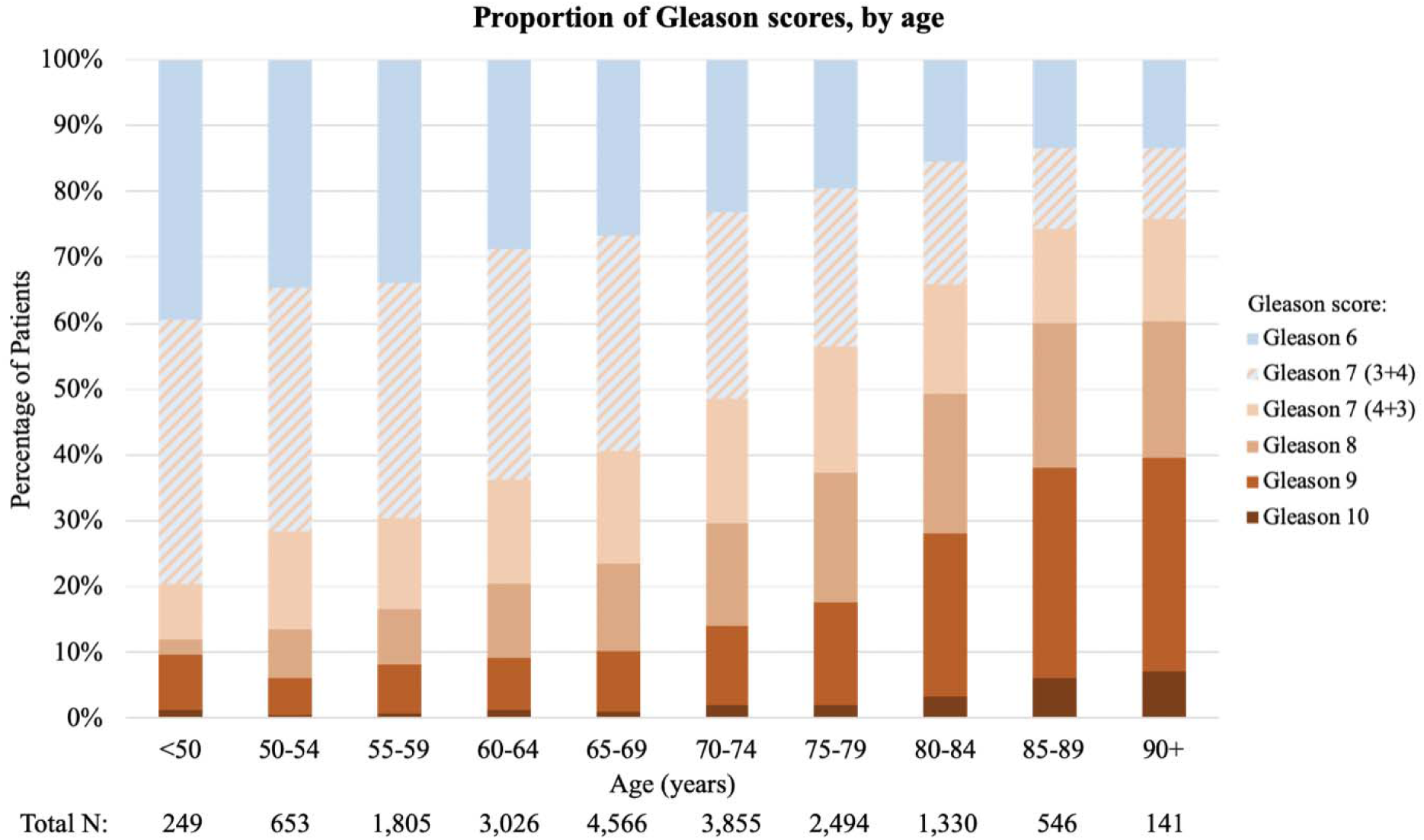
Stratification of prostate cancer patients in Norway by Gleason scores and age, 2014-2017. Patients with Gleason 7 disease were divided into Gleason 3+4 and Gleason 4+3. Total number of patients with Gleason score data available: n=18,665 (91.7% of all prostate cancer cases in Norway).

**Figure 2.**
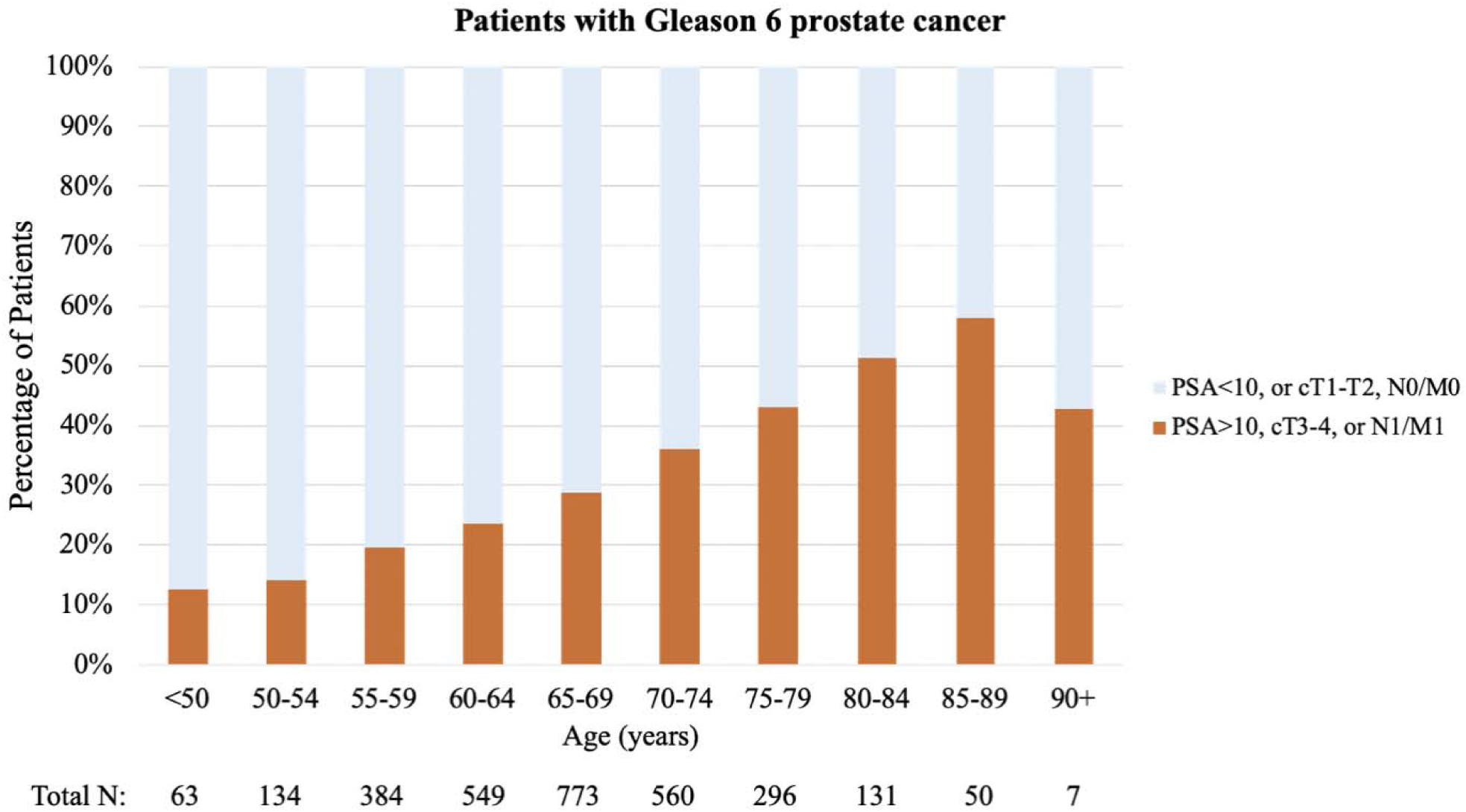
Proportion of men with Gleason 6 prostate cancer who meet one or more of the following criteria: PSA ≥10 ng/mL, clinical T3-4 stage, or N1/M1 disease, by age group, n=2,947 men.

Older patients with PCa were also more likely to have more advanced disease. The percentage of men aged 55-59, 65-69, 75-79, and 85-89 years with at least high-risk PCa was 29.3%, 39.1%, 60.4%, and 90.6%, respectively. The percentage of men with low-risk PCa across the same age groups was 24.0%, 17.9%, 10.2%, and 4.1%, respectively (p<0.001). **Figure 3** shows age-stratified proportions of cases by PCa risk group. Older men were more likely to be diagnosed with clinically-significant disease: p<0.001 when counting at least intermediate-risk as clinically-significant (the percentage of men aged 55-59, 65-69, 75-79, and 85-89 years with at least favorable-intermediate-risk disease was 76.0%, 82.1%, 89.8%, and 95.9%, respectively) and p<0.001 when counting at least unfavorable-intermediate-risk PCa as clinically-significant (the percentage of men aged 55-59, 65-69, 75-79, and 85-89 years with at least unfavorable-intermediate-risk disease was 53.3%, 61.2%, 75.8%, and 93.6%, respectively). When analyses were restricted to men aged 50-74, all of the above associations remained significant (p<0.001). Numerical proportions are listed in **eTable 2**.

**Figure 3.**
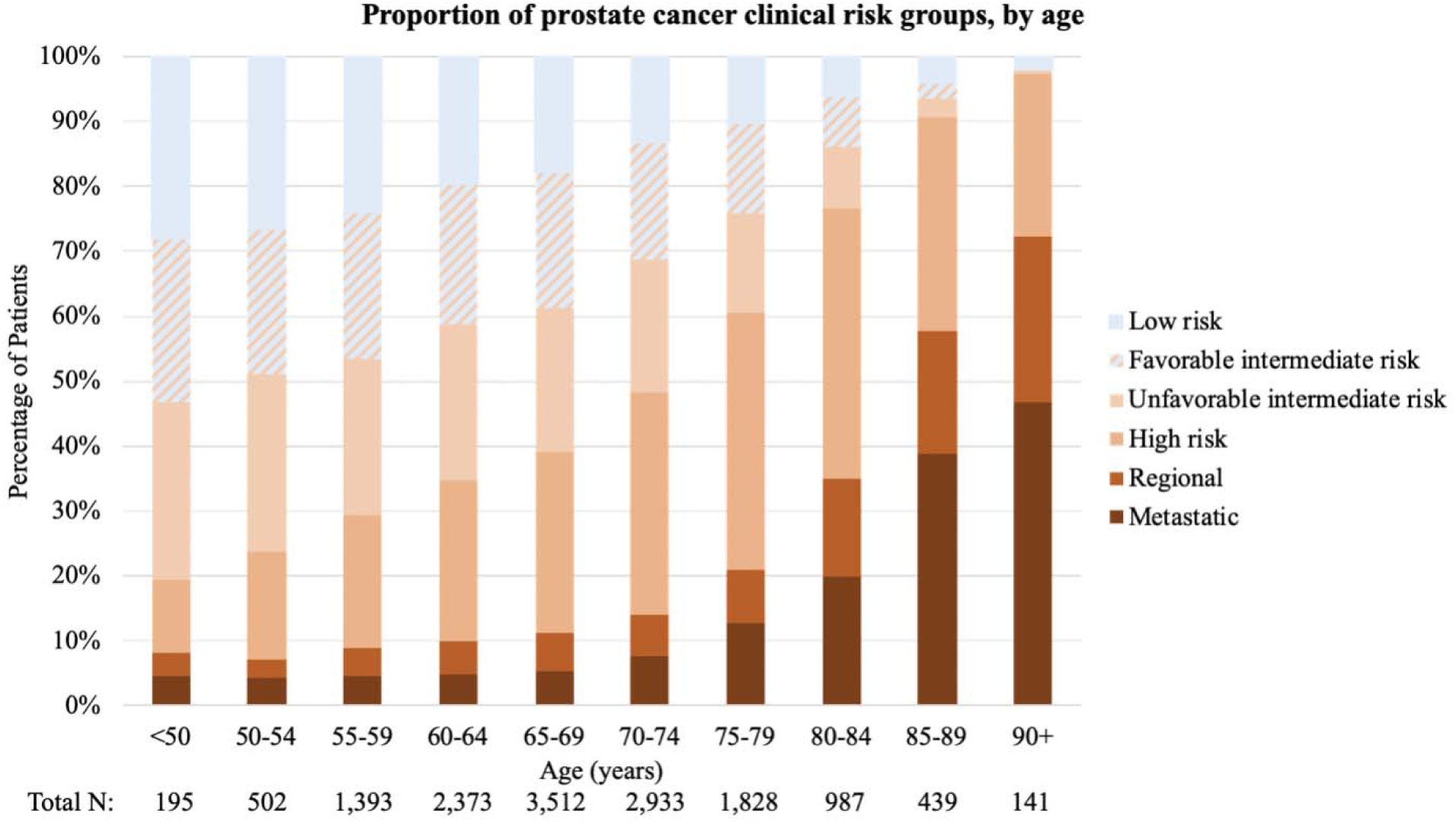
Clinical risk group stratification of prostate cancer patients in Norway, 2014-2017. Total number of patients with risk stratification data available: n=14,303 (70.3% of all prostate cancer cases in Norway).

ASIRs for risk groups are shown in **Table 1** and **Figure 4**. Older men across the Norwegian population were increasingly likely to have more advanced and clinically-significant PCa. Qualitatively, the ASIRs of PCa by risk group demonstrate an increase in incidence rates across all risk groups until ages 65-69. At ages 65-69, the rates of low and intermediate-risk cases (including both favorable and unfavorable) begin to level off or decrease. The respective maximum ASIRs for low-risk, favorable-intermediate-risk, and unfavorable-intermediate-risk PCa are: 157.1, 183.8, and 194.8, occurring at ages 65-69, 65-69, and 70-74. Meanwhile, incidence rates of high-risk disease in men over 65-69 continue to increase sharply; the ASIRs of high-risk disease surpass those of the low and intermediate-risk categories until ages 75-79—when the ASIR is 408.3—at which point they begin to decrease. The incidence of regional and metastatic disease always increases with age.

**Figure 4.**
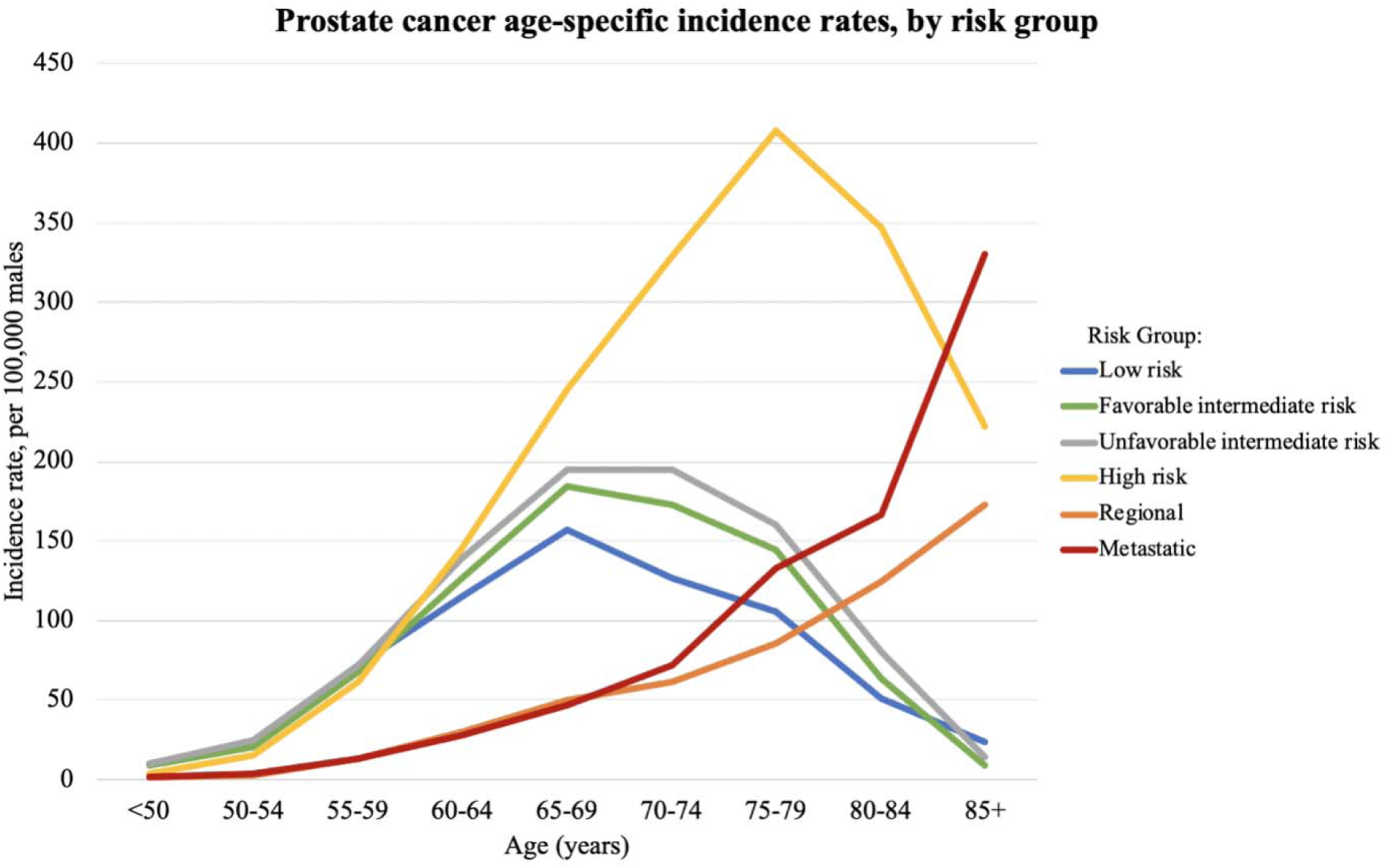
Norway prostate cancer age-specific incidence rates (per 100,000 males) by clinical risk group stratification. Intermediate-risk prostate cancer is subdivided into favorable and unfavorable risk.

## Discussion

To our knowledge, this is the first study to report PCa ASIRs by clinical risk groups. We demonstrate a strong age dependence of Gleason score and of clinical risk stratification of PCa at time of diagnosis. These findings hold true when including clinically-relevant distinctions between Gleason 3+4 and Gleason 4+3 and between favorable- and unfavorable-intermediate-risk disease^11^. Older Norwegian men are more likely to be diagnosed with higher-grade and more advanced PCa. Not only does the proportion of metastatic PCa increase with age (which could be simply due to less screening), but so does a man’s absolute risk of potentially-lethal, *localized* PCa. At age 75-79 years, the absolute incidence of high-risk, localized PCa is roughly 6 times greater than at age 55-59 years.

ASIRs for modern risk groups have important implications for screening. Older age is a striking risk factor for clinically-significant PCa in the Norwegian population, highlighting the relevance of age-specific screening decisions for individual men^1^. Rates of high-risk, localized disease increased dramatically from ages 55 to 74 (as opposed to decreasing, as in the low and intermediate-risk groups), leading to a maximum ASIR over twice as high as that of favorable intermediate-risk disease. Some guidelines discourage any PCa screening above age 70^19^, but the ASIRs shown here indicate that the absolute *incidence* of potentially-lethal disease actually *increases* in men older than 70. Healthy men aged 70-75 have a life expectancy over 10 years^20–22^, and the 10-year metastases rates for men with unfavorable-intermediate-risk and high-risk disease are high^10^. PCa remains a leading public health problem—worldwide, it is the second most common malignancy diagnosed in men and a leading cause of cancer mortality^23,24^.

The major PCa screening trials may have underestimated the potential mortality benefit of screening by including large numbers of men under 60, who had a long life-expectancy but a relatively low incidence of unfavorable-intermediate-risk or high-risk disease^16,25^. In fact, a large European trial showed a mortality benefit to screening, but on subset analysis, there was no significant benefit in men under age 65^26^. Too much focus on screening younger men (barring other risk factors) will likely tend to lead to higher relative rates of false positives and of overdiagnosis of low-risk cancer. Meanwhile, early treatment of intermediate-risk or high-risk disease could reduce PCa morbidity and mortality^5,27–29^. The present study suggests that healthy older men—those with a reasonably long life-expectancy—may be among those most likely to benefit from screening, as they are at greater risk of aggressive PCa that can cause morbidity and/or mortality if left untreated.

Our findings are consistent with prior data suggesting associations between increased age and worse PCa Gleason scores and risk groups^6–8,30^. We show these associations hold true for modern Gleason score and risk-stratification schemes. Additionally, previous work has reported population PCa stage*-* specific incidence rates and trends *over eras in time*, mainly by dividing cases into localized, regional, or distant disease^31,32^, or by using strict definitions of TNM staging^33^. The introduction of opportunistic PSA screening in Norway led to increased incidence rates of localized and regional PCa in younger men over time^31^, but no subdivision was made for localized disease that was potentially-lethal versus eligible for active surveillance. Our study shows that the favorable- and unfavorable-intermediate risk ASIRs are consistent with the overall pattern of increased locally-aggressive disease as a man ages, but the intermediate categories are closely correlated (and approximately equal to each other) across all ages. There are also continued increases in absolute rates of regional and metastatic disease in men as they age; this is an expected finding as older men are less likely to be screened for PCa^4^. Older men may also be more likely to present with metastatic disease because they were already at increased risk of having a potentially-lethal cancer, making them more prone to metastases^10^. Altogether, screening for PCa in healthy, older men (say, between 70-75 years) may allow for improved risk-stratification efforts, leading to the identification of disease while it remains potentially curable. Age could certainly be combined with other risk factors to further optimize screening strategies^1,34–37^.

Notably, Norway opted not to participate in the major European PCa screening trial^16^, citing concerns for the ethical implications of screening otherwise asymptomatic men and the overall clinical significance of the trial^38^. However, this decision was widely debated^39^, and screening remains controversial in Norway (as it is elsewhere). Norway’s national healthcare system does not recommend population-based PCa screening in asymptomatic men without a family history of PCa. Rather, shared decision-making is encouraged between physicians and patients^40^, leading to approximately 45% of Norwegian men receiving PSA testing in 2011^41^ (a rate similar to that in the USA^42^). A survey of Norwegian primary care physicians showed that a minority include PSA testing in routine tests, but 70% would order PSA testing if the patient requested, and 72% would find it “difficult” not to refer to a urologist if a patient’s PSA is elevated^41^. Physicians can use these PCa risk group ASIRs in conjunction with a patient’s other risk factors and overall health/life expectancy for maximal screening efficacy, considering the advisability and timing of possible screening (with a focus on detecting potentially-lethal disease). Knowing that older, healthy men are at the greatest risk of being diagnosed with a potentially-lethal, but potentially-curable PCa may aid physician decision-making when determining who may receive the most benefits from screening. Strategies to detect potentially-lethal disease will be most effective when the pre-test probability is high—i.e., when the underlying prevalence of clinically-significant disease is high—hence, the need to be guided by epidemiological patterns like those presented here.

Our work has limitations. We could not separate out very-low or very-high-risk cases, as PSA density information and Gleason score for each biopsy core are not registered in the Cancer Registry of Norway. These cases tend to be clinically managed similarly to low and high-risk PCa, respectively, and thus were combined with those categories^3^. Approximately 30% of the cohort could not be assigned a precise risk group due to incomplete PSA or clinical T stage information; most of these had partial data available for Gleason score and clinically-significant PCa analyses. Finally, while we obtained plausible estimates of age-specific incidence and proportion for an entire population using high-quality registry data, the direct findings are limited to Norway. Nevertheless, the patterns observed here could be similar in other western countries, a suggestion that is partially supported by prior work in the USA^6,8^.

## Conclusions

Both the proportion and absolute incidence rates of clinically-significant PCa (using modern definitions) increase with age. Notably, the absolute incidence of high-risk disease at ages 75-79 is over six times higher than that at ages 55-59. Older men are also more likely to be diagnosed with higher Gleason score. Efforts to optimize PCa screening for efficient detection of localized, potentially-lethal disease should account for this strong age dependence.

## References

1. Seibert TM, Fan CC, Wang Y, et al. Polygenic hazard score to guide screening for aggressive prostate cancer: Development and validation in large scale cohorts. BMJ. 2018;360:1–7. doi:10.1136/bmj.j5757

2. Pashayan N, Duffy SW, Chowdhury S, et al. Polygenic susceptibility to prostate and breast cancer: Implications for personalised screening. Br J Cancer. 2011;104(10):1656–1663. doi:10.1038/bjc.2011.118

3. NCCN Clinical Practice Guidelines in Oncology: Prostate Cancer version 2. 2018.

4. Parker C, Gillessen S, Heidenreich A, Horwich A, ESMO Guidelines Committee. Cancer of the prostate: ESMO Clinical Practice Guidelines for diagnosis, treatment and follow-up. Ann Oncol. 2015;26:v69–v77. doi:10.1093/annonc/mdv295

5. Hamdy FC, Donovan JL, Lane JA, et al. 10-Year Outcomes after Monitoring, Surgery, or Radiotherapy for Localized Prostate Cancer. N Engl J Med. 2016;375(15):1415–1424. doi:10.1056/NEJMoa1606220

6. Muralidhar V, Ziehr DR, Mahal BA, et al. Association between older age and increasing gleason score. Clin Genitourin Cancer. 2015;13(6):525–530e3. doi:10.1016/j.clgc.2015.05.007

7. Draisma G, Postma R, Schröder FH, Van Der Kwast TH, De Koning HJ. Gleason score, age and screening: Modeling dedifferentiation in prostate cancer. Int J Cancer. 2006;119(10):2366–2371. doi:10.1002/ijc.22158

8. Shao YH, Demissie K, Shih W, et al. Contemporary risk profile of prostate cancer in the United States. J Natl Cancer Inst. 2009;101(18):1280–1283. doi:10.1093/jnci/djp262

9. Zumsteg ZS, Spratt DE, Pei I, et al. A New Risk Classification System for Therapeutic Decision Making with Intermediate-risk Prostate Cancer Patients Undergoing Dose-escalated External-beam Radiation Therapy. Eur Urol. 2013;64(6):895–902. doi:10.1016/J.EURURO.2013.03.033

10. Spratt DE, Zhang J, Santiago-Jimenez M, et al. Development and validation of a novel integrated clinical-genomic risk group classification for localized prostate cancer. J Clin Oncol. 2018;36(6):581–590. doi:10.1200/JCO.2017.74.2940

11. Edge SB, Byrd DR, Compton CC, Fritz AG, Greene FL, Trotti A. AJCC Cancer Staging Manual, Eighth Edition.; 2017. https://www.springer.com/us/book/9783319406176. Accessed August 7, 2018.

12. Harvei S, Tretli S, Langmark F. Quality of Prostate Cancer Data in the Cancer Registry of Norway. Eur J Cancer. 1996;32(1):104–110. doi:10.1016/0959-8049(95)00501-3

13. Larsen IK, Småstuen M, Johannesen TB, et al. Data quality at the Cancer Registry of Norway: An overview of comparability, completeness, validity and timeliness. Eur J Cancer. 2009;45(7):1218–1231. doi:10.1016/j.ejca.2008.10.037

14. Epstein JI, Egevad L, Amin MB, Delahunt B, Srigley JR, Humphrey PA.The 2014 International Society of Urological Pathology (ISUP) Consensus Conference on Gleason Grading of Prostatic Carcinoma. Am J Surg Pathol. 2015;40(2):1. doi:10.1097/PAS.0000000000000530

15. R Core Team. R: A language and environment for statistical computing. In: Vienna, Austria: R Foundation for Statistical Computing.; 2015.

16. Schröder FH, Hugosson J, Roobol MJ, et al. Screening and Prostate-Cancer Mortality in a Randomized European Study. N Engl J Med. 2009;360(13):1320–1328. doi:10.1016/j.eeh.2004.05.002

17. Engholm G, Ferlay J, Christensen N, et al. NORDCAN - A Nordic tool for cancer information, planning, quality control and research. Acta Oncol (Madr). 2010;49(5):725–736. doi:10.3109/02841861003782017

18. Engholm G, Ferlay J, Christensen N, et al. Cancer Incidence, Mortality, Prevalence and Survival in the Nordic Countries, Version 8.1 (28.06.2018). Association of the Nordic Cancer Registries. Danish Cancer Society. http://www.ancr.nu. Accessed September 18, 2018.

19. Grossman DC, Curry SJ, Owens DK, et al. Screening for prostate cancer: US Preventive servicestaskforcerecommendation statement. JAMA - J Am Med Assoc. 2018;319(18):1901–1913. doi:10.1001/jama.2018.3710

20. Social Security Administration. Actuarial Life Table 2014. https://www.ssa.gov/OACT/STATS/table4c6.html. Accessed March 27, 2019.

21. Statistisk sentralbyrå (Statistics Norway). Life Expectancy - Remaining Years for Males and Females at Selected Ages. Statistisk sentralbyra https://www.ssb.no/en/befolkning/statistikker/dode/aar. Accessed March 27, 2019.

22. NCCN. Older Adult Oncology v2.2018. NCCN. 2018.

23. Bray F, Ferlay J, Soerjomataram I, Siegel RL, Torre LA, Jemal A. Global cancer statistics 2018: GLOBOCAN estimates of incidence and mortality worldwide for 36 cancers in 185 countries. CA Cancer J Clin. 2018;68(6):394–424. doi:10.3322/caac.21492

24. Ferlay J, Colombet M, Soerjomataram I, et al. Cancer incidence and mortality patterns in Europe: Estimates for 40 countries and 25 major cancers in 2018. Eur J Cancer. 2018;103:356–387. doi:10.1016/J.EJCA.2018.07.005

25. Andriole GL, Crawford ED, Grubb RL, et al. Mortality Results from a Randomized Prostate-Cancer Screening Trial. N Engl J Med. 2009;360(13):1310–1319. doi:10.1056/NEJMoa0810696

26. Schröder FH, Hugosson J, Roobol MJ, et al. Screening and prostate cancer mortality: Results of the European Randomised Study of Screening for Prostate Cancer (ERSPC) at 13 years of follow-up. Lancet. 2014;384(9959):2027–2035. doi:10.1016/S0140-6736(14)60525-0

27. Bill-Axelson A, Holmberg L, Garmo H, et al. Radical Prostatectomy or Watchful Waiting in Prostate Cancer — 29-Year Follow-up. N Engl J Med. 2018;379(24):2319–2329. doi:10.1056/NEJMoa1807801

28. Bolla M, de Reijke TM, Van Tienhoven G, et al. Duration of Androgen Suppression in the Treatment of Prostate Cancer. N Engl J Med. 2009;360(24):2516–2527. doi:10.1056/NEJMoa0810095

29. Jones CU, Hunt D, McGowan DG, et al. Radiotherapy and Short-Term Androgen Deprivation for Localized Prostate Cancer. N Engl J Med. 2011;365(2):107–118. doi:10.1056/NEJMoa1012348

30. Alibhai SMH, Krahn MD, Fleshner NE, Cohen MM, Tomlinson GA, Naglie G. The association between patient age and prostate cancer stage and grade at diagnosis. BJU Int. 2004;94(3):303–306. doi:10.1111/j.1464-410X.2004.04883.x

31. Møller MH, Kristiansen IS, Beisland C, Rørvik J, Støvring H. Trends in stage-specific incidence of prostate cancer in Norway, 1980–2010: a population-based study. BJU Int. 2016;118(4):547–555. doi:10.1111/bju.13364

32. Li J, Siegel DA, King JB. Stage-specific incidence rates and trends of prostate cancer by age, race, and ethnicity, United States, 2004–2014. Ann Epidemiol. 2018;28(5):328–330. doi:10.1016/j.annepidem.2018.03.001

33. Cremers RGHM, Karim-Kos HE, Houterman S, et al. Prostate cancer: Trends in incidence, survival and mortality in the Netherlands, 1989-2006. Eur J Cancer. 2010;46(11):2077–2087. doi:10.1016/j.ejca.2010.03.040

34. Vickers AJ, Ulmert D, Sjoberg DD, et al. Strategy for detection of prostate cancer based on relation between prostate specific antigen at age 40-55 and long term risk of metastasis: casecontrol study. BMJ. 2013;346:f2023. doi:10.1136/bmj.f2023

35. Ankerst DP, Straubinger J, Selig K, et al. A Contemporary Prostate Biopsy Risk Calculator Based on Multiple Heterogeneous Cohorts. Eur Urol. 2018;74(2):197–203. doi:10.1016/j.eururo.2018.05.003

36. Ankerst DP, Hoefler J, Bock S, et al. Prostate cancer prevention trial risk calculator 2.0 for the prediction of low- vs high-grade prostate cancer. Urology. 2014;83(6):1362–1367. doi:10.1016/j.urology.2014.02.035

37. Huynh-Le M-P, Fan CC, Parsons JK, et al. A genetic risk score to personalize prostate cancer screening, applied to population data. J Clin Oncol. 2019;37(7_suppl):181–181. doi:10.1200/jco.2019.37.7_suppl.181

38. Fosså SD, Eri LM, Skovlund E, Tveter K, Vatten L, Norwegian Urological Cancer Group. No randomised trial of prostate cancer screening in Norway. Lancet Oncol. 2001;2(12):741–745. doi:10.1016/S1470-2045(01)00588-5

39. Schröder FH, De Koning HJ. The Norwegian decision on screening for prostate cancer: A response. Lancet Oncol. 2001;2(12):746–749. doi:10.1016/S1470-2045(01)00589-7

40. Norwegian Directorate of Health. Screening for prostatakreft i den generelle befolkningen - Nasjonalt handlingsprogram med retningslinjer for diagnostikk, behandling og oppfølging av prostatakreft. https://www.helsebiblioteket.no/retningslinjer/prostatakreft/3-screening-og-tidlig-påvisning/3.1-screening-for-prostatakreft. Accessed February 2, 2019.

41. Breidablik HJ, Meland E, Aakre KM, Førde OH. PSA measurement and prostate cancer – overdiagnosis and overtreatment? Tidsskr Nor Legeforen. 2013;16(133):1711–1716.

42. Li J, Berkowitz Z, Hall IJ. Decrease in Prostate Cancer Testing Following the US Preventive Services Task Force (USPSTF) Recommendations. J Am Board Fam Med. 2015;28(4):491–493. doi:10.3122/jabfm.2015.04.150062

