## Supplemental Figure 1 and Tables 1-2 for "Age dependence of modern clinical risk groups for localized prostate cancer – a population-based study"

**Supplemental eFigure 1**. Proportion of men with Gleason 7 (3+4) prostate cancer who meet one or more of the following criteria: PSA ≥10 ng/mL, clinical T3-4 stage, or N1/M1 disease, by age group (Panel A: n=3,946 men). Panel B shows the proportion of men with Gleason 6 or 7 (3+4) disease who meet the same criteria (n=6,893 men).

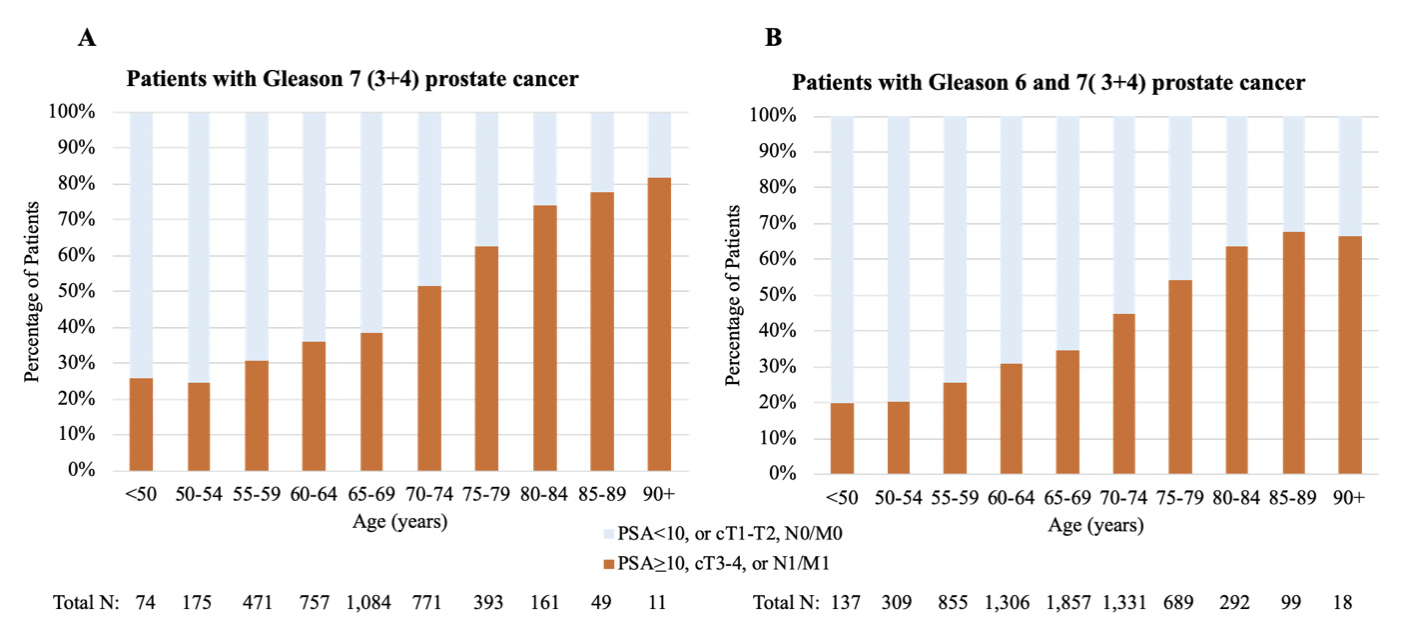

**Supplemental eTable 1**. Proportion of men with prostate cancer in Norway by Gleason scores and age, 2014-2017. Patients with Gleason 7 disease were divided into Gleason 3+4 and Gleason 4+3. Total number of patients with Gleason score data available: n=18,665 (91.7% of all prostate cancer cases in Norway).

|  | **Proportion of men with each Gleason score** | | | | | |
| --- | --- | --- | --- | --- | --- | --- |
| **Age (years)** | **6** | **7 (3+4)** | **7 (4+3)** | **8** | **9** | **10** |
| **<50** | 39.4 | 40.2 | 8.4 | 2.4 | 8.4 | 1.2 |
| **50-54** | 34.5 | 37.1 | 15.0 | 7.3 | 5.5 | 0.6 |
| **55-59** | 33.8 | 35.7 | 14.0 | 8.4 | 7.3 | 0.8 |
| **60-64** | 28.8 | 34.9 | 15.8 | 11.2 | 8.1 | 1.2 |
| **65-69** | 26.8 | 32.5 | 17.3 | 13.2 | 9.1 | 1.1 |
| **70-74** | 23.2 | 28.2 | 19.1 | 15.5 | 12 | 2.0 |
| **75-79** | 19.5 | 24.0 | 19.3 | 19.7 | 15.4 | 2.1 |
| **80-84** | 15.4 | 18.8 | 16.5 | 21.2 | 24.8 | 3.3 |
| **85-89** | 13.4 | 12.4 | 14.3 | 21.8 | 32.1 | 6.0 |
| **90+** | 13.5 | 10.6 | 15.6 | 20.6 | 32.6 | 7.1 |

**Supplemental eTable 2**. Proportion of men with prostate cancer in Norway by modern risk group and age, 2014-2017. Total number of patients with risk stratification data available: n=14,303 (70.3% of all prostate cancer cases in Norway).

|  | **Proportion of men in each prostate cancer risk group** | | | | | |
| --- | --- | --- | --- | --- | --- | --- |
| **Age (years)** | **Low-risk** | **Favorable intermediate-risk** | **Unfavorable Intermediate-risk** | **High-risk** | **Regional** | **Metastatic** |
| **<50** | 28.2 | 25.1 | 27.2 | 11.3 | 3.6 | 4.6 |
| **50-54** | 26.5 | 22.5 | 27.3 | 16.5 | 3.0 | 4.2 |
| **55-59** | 24.0 | 22.7 | 24.0 | 20.4 | 4.4 | 4.5 |
| **60-64** | 19.7 | 21.6 | 23.9 | 24.9 | 5.1 | 4.8 |
| **65-69** | 17.9 | 20.9 | 22.1 | 28.0 | 5.7 | 5.4 |
| **70-74** | 13.3 | 18.1 | 20.3 | 34.3 | 6.4 | 7.6 |
| **75-79** | 10.2 | 14 | 15.4 | 39.4 | 8.2 | 12.8 |
| **80-84** | 6.2 | 7.6 | 9.6 | 41.6 | 15.0 | 20.0 |
| **85-89** | 4.1 | 2.3 | 3.0 | 32.8 | 19.1 | 38.7 |
| **90+** | 2.1 | 0 | 0.7 | 24.8 | 25.6 | 46.8 |
